# Sirt3 enhanced Bufalin sensitivity of colorectal cancer cells through P53- dependent pathway of apoptosis

**DOI:** 10.1101/2022.08.16.504176

**Authors:** Lu Li, Cui Liu, Xinqing Zhang, Xiao Meng, Yaxiao Geng, Zhigang Shi, Yongzhi Zhang

**Author notes:** Correspondence to: Lu Li.

## Abstract

Sirt3, one of class III histone deacetylase, is mainly localized in mitochondria and plays a significant role in the control of the metabolic activity, senescence and death [1]. Recently, Sirt3 emerged as a novel member of anticancer. However, the role of Sirt3 in colorectal cancer (CRC) has never been explained exactly. In this study, we found that sirt3 is down-regulated after Bufalin treatment. We also found that AC-P53 is up-regulated which induces Bax translocation to mitochondrion and open the mitochondrial permeability transition (mPTP) pores result in the release of cytochrome C lead to the activation of caspase-dependent apoptosis pathway. Collectively, our data suggests that Sirt3 may play an important role in CRC development and progression and may be a promising therapeutic target for CRC.

## Introduction

Cancer is a major public health problem worldwide and is the second leading cause of death in the United Stat1es [2]. The aim of apoptosis is to eliminate damaged or abnormal cells: Cancer cells, however, cancer cells adopt various strategies to override apoptosis for survival [3].It is crucial that to understand the mechanisms of apoptosis for preventing and curing cancer.

Bufalin is an active compound in the traditional Chinese medicine Chan Su, which has been recently reported as a potent anticancer drug in various cancers, such as hepatocellular carcinoma, CRC and Lung Cancer [4–6].However, the genetic mechanism underlying has not been elucidated yet. The purpose of our study was to investigate whether mitochondria‑mediated signaling pathways trigger the process of apoptosis in the CRC cell lines.

Sirt3, a member of the silent information regulator 2 (SIR2) family, is a NAD+-dependent protein deacetylase [7]. SIRT3 has received much attention for its role in cancer genetics, aging, neurodegenerative disease, and stress resistance [8]. SIRT3 is mainly located within mitochondria and had an important role in mitochondrial homeostasis [9]. The tumor suppressor p53 is an essential and key transcription factor that undergoes an abundance of post-translational modifications for its regulation and activation, including phosphorylation, acetylation, and methylation. Acetylation of p53 is an important reversible enzymatic process and is indispensable essential for p53 transcriptional activity. p53 was the first non-histone protein shown to be acetylated by histone acetyl transferases[10]. And it has reported that sirt3 contributes to the acetylation or deacetylation of P53 [11].As we all know, P53 promotes the translocation of Bax to mitochondria and the release of Cytochrome C[12].What we want to know is that if is the acetylated of P53 promotes the translocation of Bax to mitochondria. And our results confirmed our hypothesis.

## Results

### Effect of Bufalin on cell viability and cell cycle

Effect of Bufalin on cell viability, HCT-116 and HCT-116 Bax−/− cells, were tested. MTT shows that Bufalin decreased cell viability both in a dose – and time – dependent manner (Figure 2A). Flow cytometry showed that the number of SubG1 cells was significantly increased in those Treated with 0.05, 0.5μM Bufalin in a dose-dependence manner (Figure 2B).

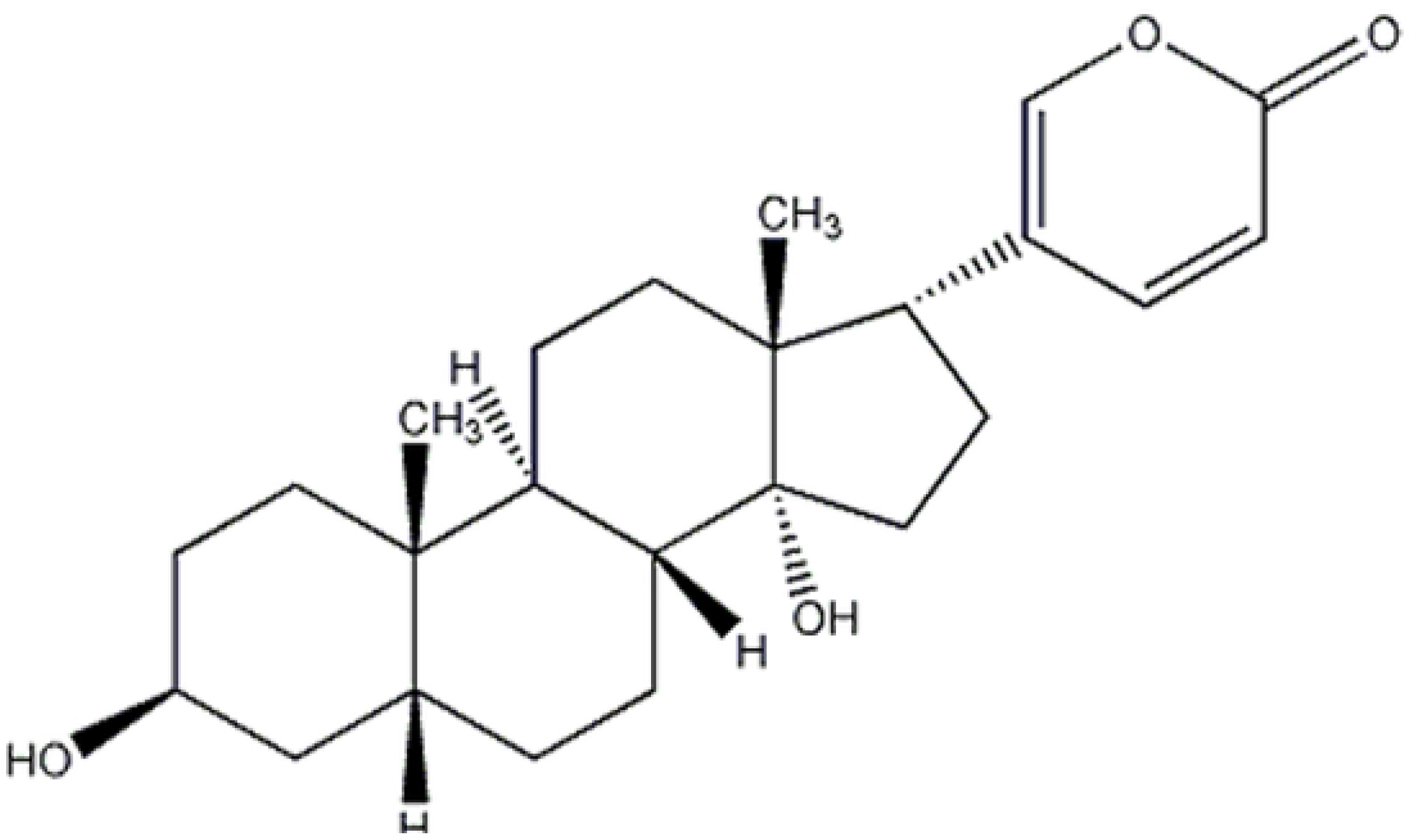

**Figure2.**
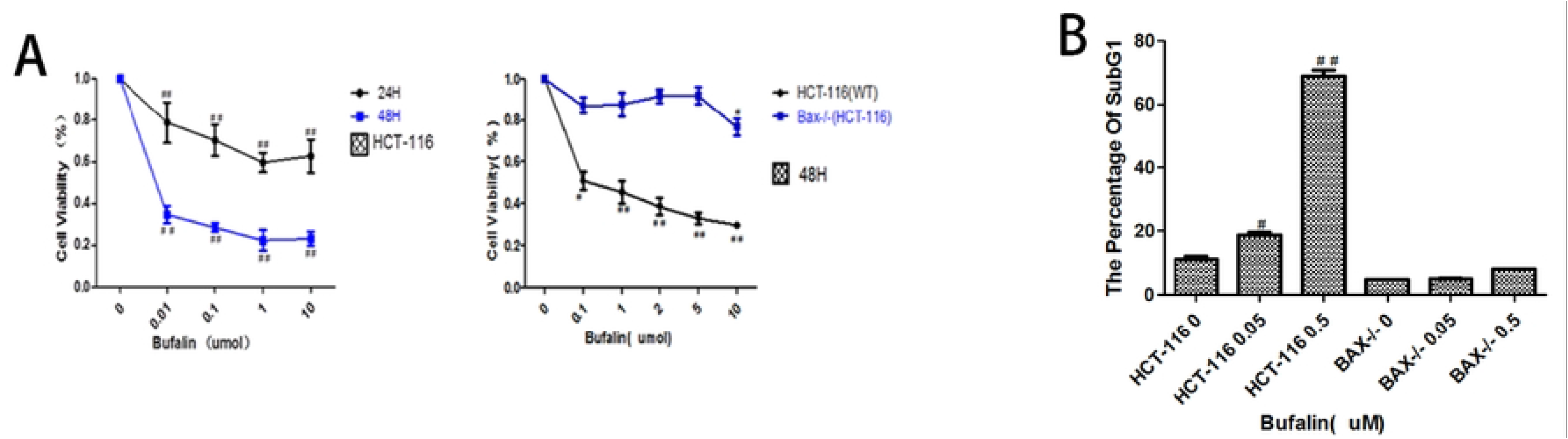
Effect of Bufalin on cell viability and cell cycle.(A) MTT shows that Bufalin decreased cell viability both in a dose-and time – dependent manner. (B) Flow cytometry showed that the number of SubG1 cells was significantly increased in a dose-dependence manner.( #P< 0.05, ##P<0.01)

### Bufalin induces HCT-116 cells apoptosis and is BAX-depedent

To adress which cell death subroutine was answerable to the lowered viability. Hoechst 33342 staining and Annexin V/7-AAD double staining were carried out, and the result showed that most of the cell death induced by Bufalin can be classified as apoptosis in HCT-116 and is BAX-depedent. Activation of caspases is a biochemical feature of apoptosis[16]. Immunoblotting assessment showed that caspase 3 was cleaved by Bufalin. The cleaved caspase 3 were increased by Bufalin in a dose-dependent manner. The cleavage of poly (ADP) ribose polymerase (PARP),a caspase-3/7 substrate[17], was also increased by Bufalin treatment (Figure 3C). These morphological and proteins changes suggest that the cell death caused by Bufalin is apoptosis and is BAX-depedent. These phenomenons of apoptosis were obviously found when Bufalin treated on HCT-116 cells, but on HCT-116 Bax−/− cells did not found, So we come to the conclusion that Bufalin induces HCT-116 cells apoptosis and this manner of death is BAX-dependent.

**Figure 3:**
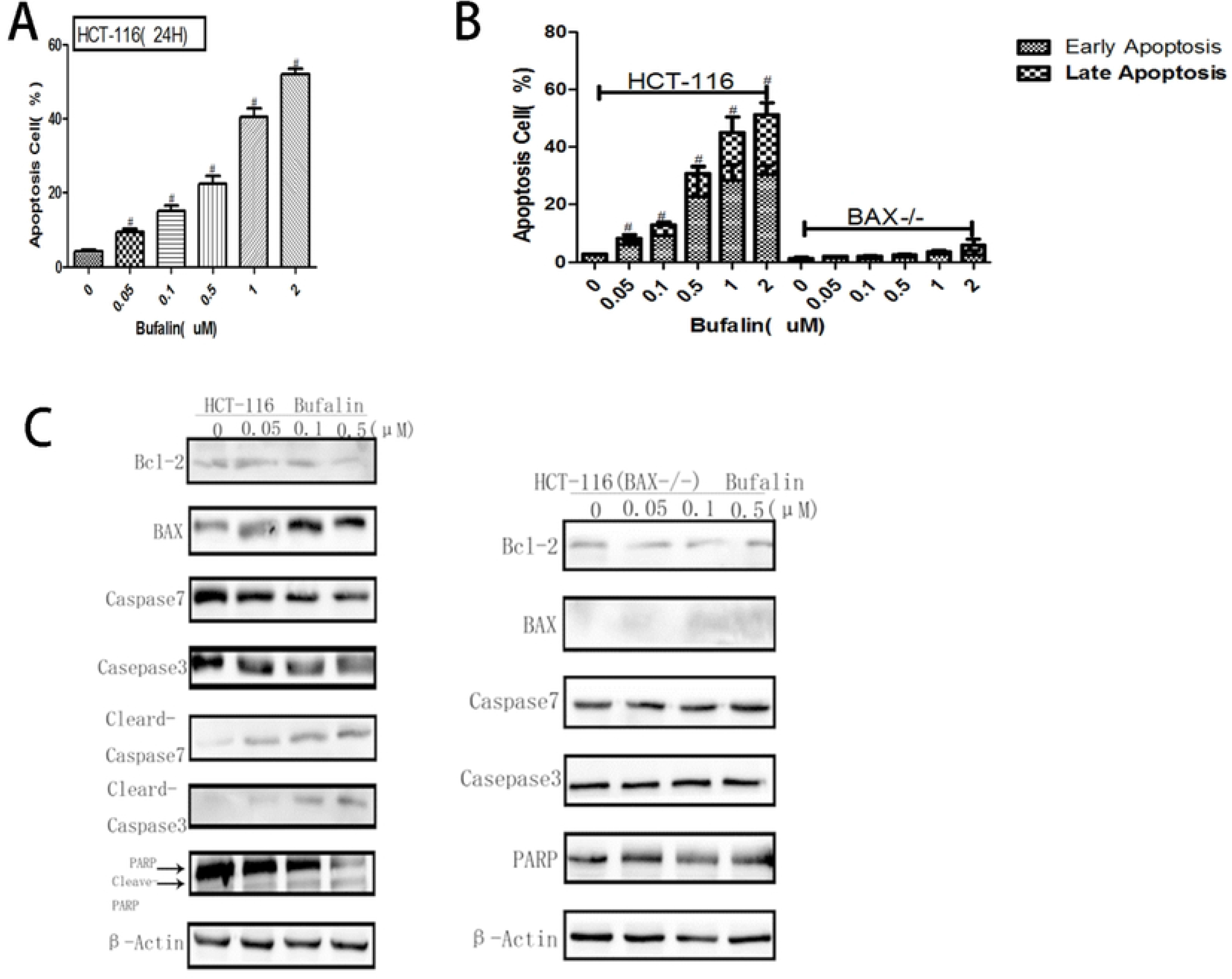
Bufalin induces HCT-116 cells apoptosis and is BAX-depedent. (**A**)Cells were exposured to Hoechst33342 (5 μg/ml) and subjected to analysis of apoptosis rate (n = 3). (**B**)PE-annexin V/7-AAD exposed on the cell surface was measured by flow cytometric analysis (n = 3). Data came from three separate experiments and was expressed as the mean ± SD. (**C**)Total cell lysates were prepared for Immunoblotting analysis of the apoptosis relative proteins (n = 3).

### Bufalin increases releasement of reactive oxygenspecies (ROS) and cytochrome C in HCT-116 cells

ROS induces mitochondria dysfunction to release proapoptotic molecules, and results in the activation of caspase-9 and caspase-3 during apoptosis [18–20]. The releases of cytochrome C illustrates that Bufalin can change the mitochondrial membrane permeability transition pore (mPTP).So, Bufalin induces mitochondrial apoptotic pathways in HCT-116 cells.

### Bufalin down-regulated the expression of sirt3 in CRC cells and this result also illustrated that Bax, P53 is the downstream target genes of sirt3

As sirt3 resides inside mitochondria and governs mitochondrial ROS generation, energy metabolism, biogenesis and apoptosis [21, 22].P53 have been reported as sirt1 down targets [23]. Sirt3 may be identified as tumor suppressor due to it can inhibit mitochondrial ROS generation as ROS are involved in both the initiation and promotion of tumorigenesis [24].Bufalin induces ROS generation (Figure 4A), so we detected the expression of sirt3 in mRNA and protein after Buffalin treatment(Figure 5A–C). Bufalin down-regulated the expresson of sit3 in CRC cells, Bax and P53 deficiency or expression has no effect on the expression of sirt3.

**Figure 4:**
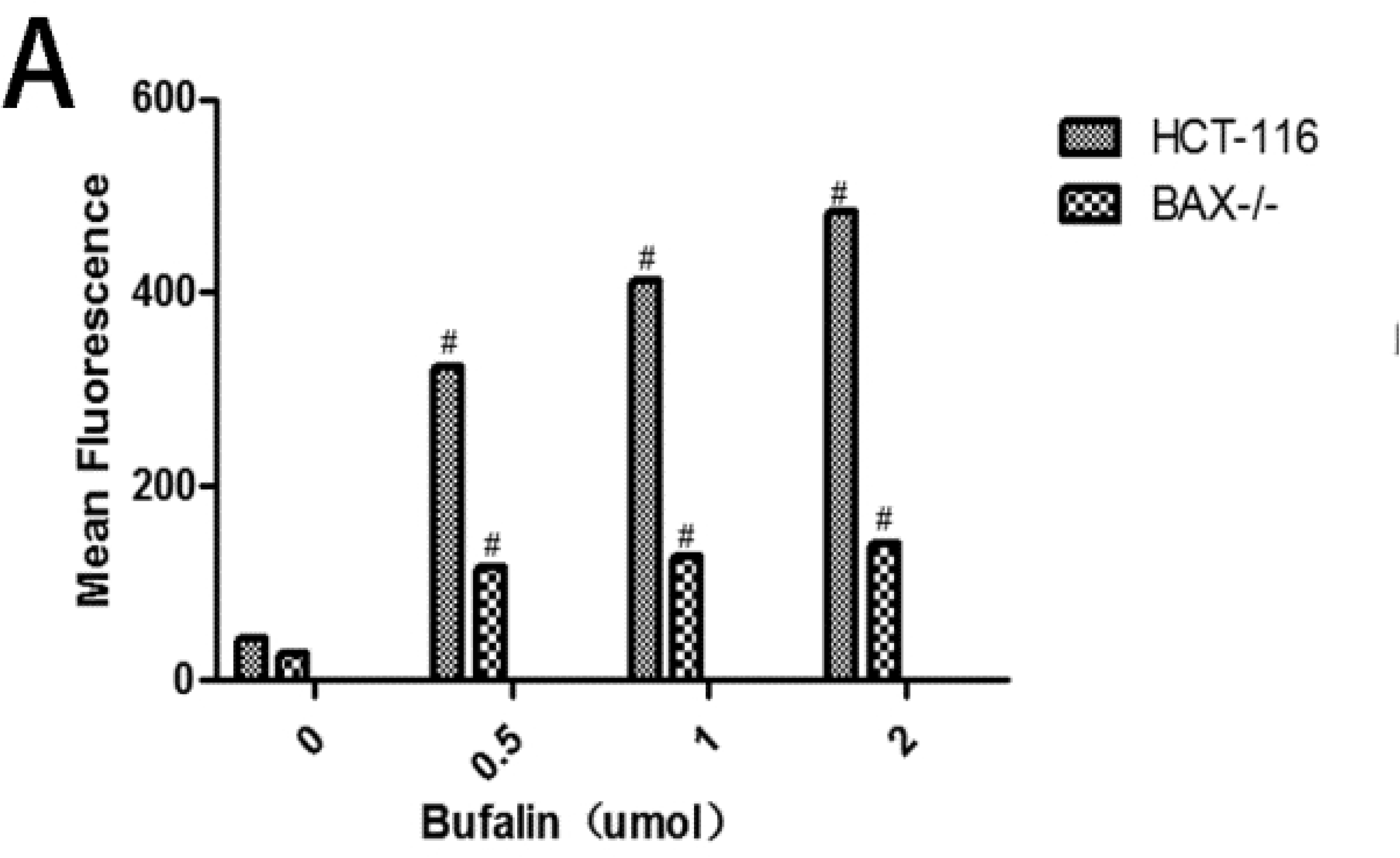
Bufalin increases releasement of reactive oxygenspecies (ROS) and cytochrome C in HCT-116 cells. (**A**) Bufalin increases ROS production in HCT-116 cells. Data came from three separate experiments and was expressed as the mean ± SD(n=3).^#^ *p* < 0.05, ^##^ *p* < 0.01

**Figure 5:**
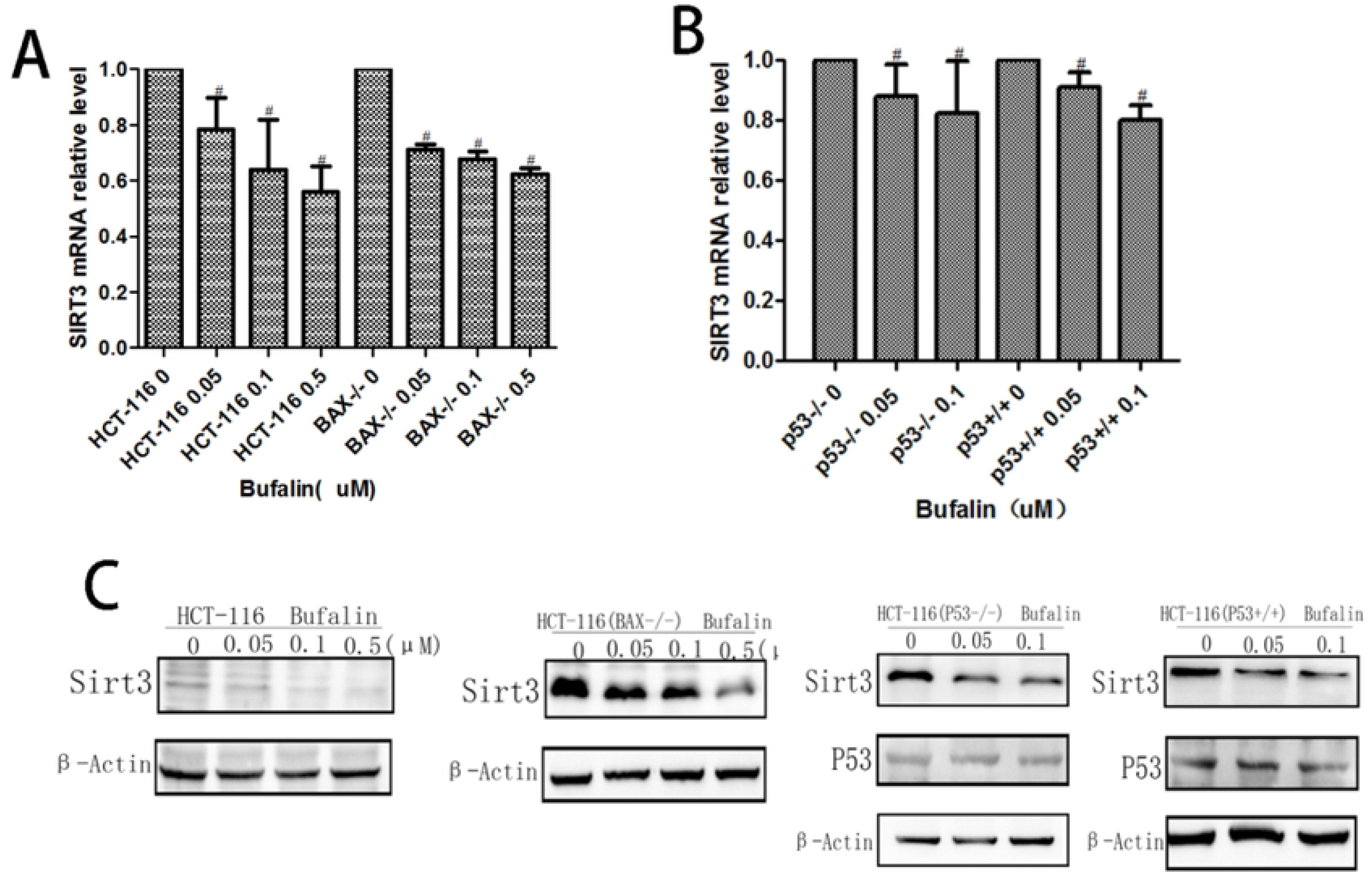
Bufalin down-regulated the expression of sirt3 in CRC cells and this result also illustrated that Bax, P53 is the downstream target genes of sirt3. (A) and (B) HCT116,HCT116 Bax−/−, HCT116 P53−/− and HCT116 P53+/+ cells were treated with Bufalin for 12 h. RNA was extracted and qRT-PCR were performed to analyze sitr3 mRNA relative level.(C) Ptotein expression of sirt3 in CRC cells was detected by Western blotting.

### Bufalin down-regulated the expression of sirt3 in HCT – 116 cells, inhibited the interaction of sirt3 and P53 and increased AC-P53 expression

On the other hand, Sirt3 is believed to be the major protein deacetylase within the mitochondrion [25, 26], and we observed that Bufalin can make Sirt3 and P53 down-regulation, Ac-p53 up-regulation, and P-p53 unchanged(Figure 6A). To determine if the specific association between Sirt3 and P53 is required for the induction of intrinsic apoptosis, coimmunoprecipitation was performed in HCT116 cells. A strong interaction between Sirt3 and P53 was found in HCT116 under apoptotic condition when proteins were immunoprecipitated with Sirt3 or P53 antibody, and the interaction is dose-dependent decrease of Bufalin (Figure 6B and C).We also used Sirt3 siRNA to confirm this result. When Sirt3 is knocked down, the same result with Bufalin (Figure 6D).

**Figure 6:**
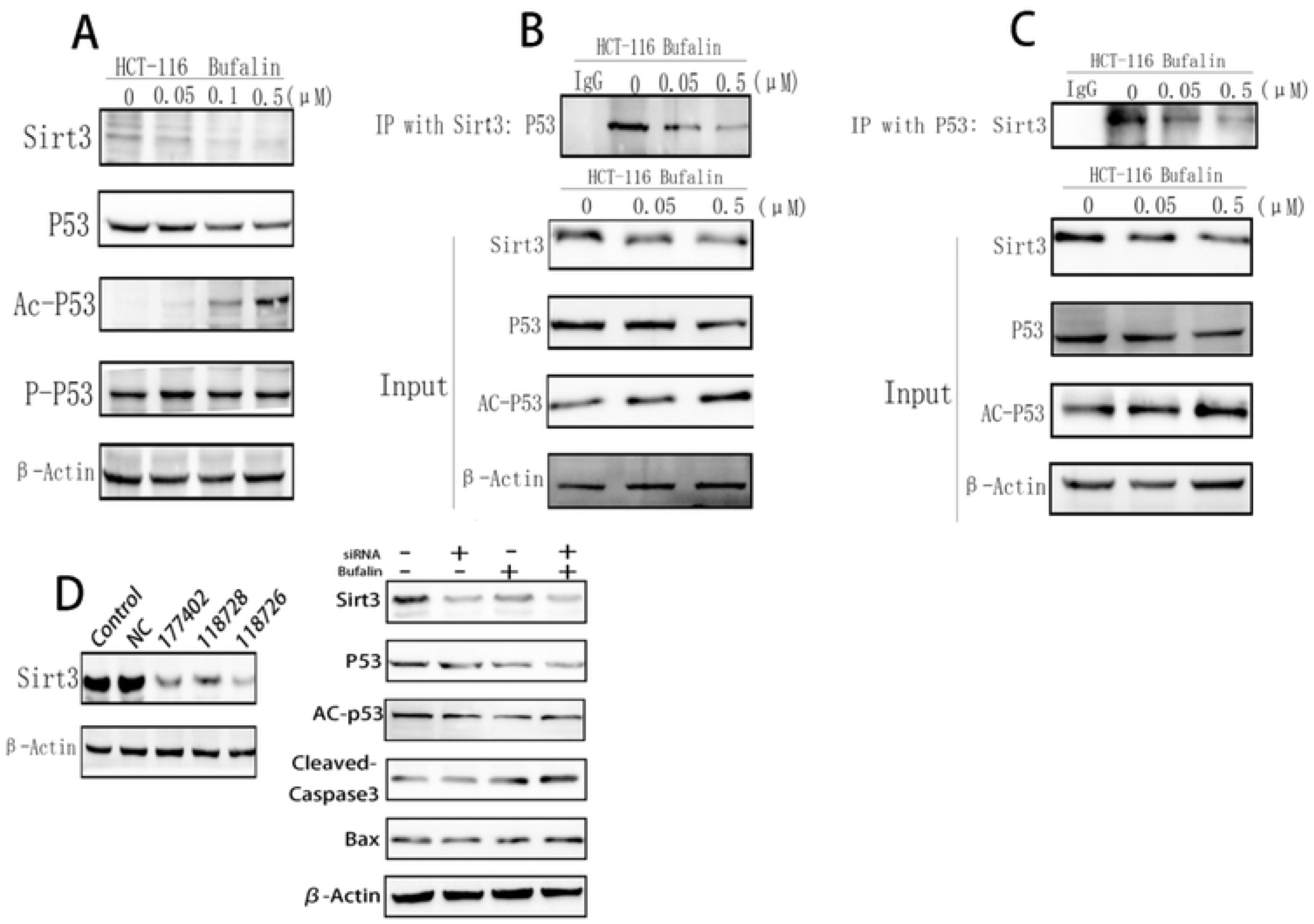
Bufalin down-regulated the expression of sirt3 in HCT – 116 cells, inhibited the interaction of sirt3 and P53 and increased AC-P53 expression. (A) HCT116 cells were treated with Bufalin for 12 h. Total cell lysates were prepared for Western blotting analysis of the Sirt3、P53、Ac-p53(Lys-382) and P-p53 proteins (n = 3). (B) and (C) IP was performed with an anti-Sirt3 or anti-P53 antibody. Co-IP Sirt3 and P53 were detected by western blotting (n = 3). (D) HCT116 cells were transfected with Sirt3 siRNA, and the cells were then treated with Bufalin for 12 h. Western blotting was performed with Sirt3、Bax、Cleaved-Caspase3、P53、Ac-p53 and P-p53 antibodies(n = 3).

### AC-p53 is key factor of Bufalin induced intrinsic apoptosis in HCT-116 cells, which promotes the translocation of Bax to mitochondrial outer membrane

It has reported that Metformin-induced decreases in Ac-p53 [27], we observed that Metformin indeed decreases the expression of Ac-p53 and has no effect on apoptosis in HCT-116 cell (Figure 7A–C).When HCT-116 treated with Metformin(10mM)and Bufalin(0.1uM),the expression of Cleaved-Caspase 3 and Cleaved-Parp is declined, but the Bax is rised (Figure 7C). So, we can come to a conclusion AC-p53 is key factor of Bufalin induced intrinsic apoptosis in HCT-116 cells. But also, by Confocal, we observed that AC-p53 promotes the translocation of Bax to mitochondrial outer membrane (Figure 7D).

**Figure 7:** AC-p53 is key factor of Bufalin induced intrinsic apoptosis in HCT-116 cells, which promotes the translocation of Bax to mitochondrial outer membrane. (A) HCT116 cells were treated with Metformin for 12 h. Total cell lysates were prepared for Western blotting analysis of Ac-p53 (Lys-382) (n = 3). (B) and (C) Western blotting was performed with antibodies of Ac-p53 (Lys-382)、Bax、Cleaved-Caspase3、P53、and P-p53(n = 3). (D) Confocal immunofluorescence of Bax (red, anti-Bax) in the HCT-116 cells that were loaded with Mito-green and DAPI. 1000× for all, scale bar = 10 μm. (n = 3).

## Discussion

Bufalin is an active compound in the traditional Chinese medicine Chan Su, is studied extensively for its multitude of pharmacological actions including anticancer properties. Our experimental data indicated that Bufalin mediated dose dependent inhibitory action on HCT-116 cells proliferation was due to AC-p53 up-regulation. It has been reported that P53 is a tumor suppressor and regulates processes like cell cycle, apoptosis, autophagy and DNA damage repair [28]. Transcriptional activity of p53 is also controlled by post-translational modification like phosphorylation and acetylation. Interestingly, acetylation is crucial for the progress of apoptosis [29]. Acetyl-p53 is known to play determining role in transcription-independent pathway of Bax-mediated apoptosis [30, 31]. Here, we investigated the mechanisms underlying the effect of Bufalin on the regulation of p53 and p53-related apoptosis pathways. We found that the Bufalin inhibited the Sirt3-mediated p53 deacetylation in HCT-116 cells. Furthermore, Bufalin had a stronger inhibitory effect on apoptosis in HCT-116 cells than in HCT-116 cells Bax−/− cells (Figure 3).

Our studies suggested that sirt3-mediated deacetylation played an important role in Bufalin-induced apoptosis. Sirt1 is the homologous protein of Sirt3, and negatively regulates the tumor suppressor P53 [32]. Therefore, we come to a hypothesis that Sirt3 may play a similar role as Sirt1. Our results showed that P53 could bind with Sirt3, and Bufalin had effect on their interaction in a dose-dependent way (Figure 6B–C). However, Bufalin decreased Sirt3 protein level in HCT-116 cells (Figure 6A). We found that Bufalin enhanced P53 acetylation via a different mechanism related with Sirt3 instead of influencing the direct action between P53 and Sirt3.Furthermore,we also found that there has no effect on the expression of sirt3 in HCT-116 Bax−/−、HCT-116 P53−/−、HCT-116 P53+/+ cells treated with Bufalin of various concentrations (Figure 5). This illustrated that Bax and P53 is the downstream target genes of Sirt3.

Acetylation is an important mechanism of regulation of P53 function, Sirt3 played a critical role in the acetylation of P53 induced by Bufalin. As most studies support a mitochondrial localization and deacetylase activity for Sirt3[33], others report that both forms of SIRT3 are enzymatically active[34]. Prof. Reinberg Iwahara et al. report that SIRT3 is capable of histone deacetylase (HDAC) activity [35]. Our findings that Bufalin decreased Sirt3 induced the deacetylation of P53 suggest that Bufalin up-regulates Ac-p53 to induce translocation of Bax to promote the progress of apoptosis.

## Materials and methods

### Materials

Bufalin (Fig.1), dimethyl sulfoxide (DMSO), and thiazolyl blue tetrazolium bromide (MTT) were obtained from Sigma Chemical Co (St. Louis, MO, USA). RPMI 1640, FBS, and antibiotics were obtained from Invitrogen (Gibco, Carlsbad, CA, USA). Apoptosis Detection Kit was purchased from BD Biosciences (USA). Antibodies were bought from Cell Signaling Technology.

### Cell culture

Human Colon cell lines HCT-116、HCT-116 Bax−/− 、HCT-116 P53−/−、HCT-116 P53+/+ were purchased from American Type Culture Collection (ATCC, Rockville,MD). HCT-116 were cultured in RPMI-1640(Gibco), HCT-116 P53−/− 、 HCT-116 P53+/+ cultured in DMEM (Gibco),and HCT-116 Bax−/− cultured in McCoy’s 5A (Gibco).All cell lines were carried out in 10% (v/v) fetal bovine serum (Invitrogen) and 1% (v/v) penicillin– streptomycin (Invitrogen) in 5%CO2 at 37°C. Transient transfection of sirt3 siRNA was carried out using Neofect Transfection Reagent following the instructions of the manufacturer.

### Cell viability assay

The effect of Bufalin on the viability of HCT-116、HCT-116 Bax−/− cells was examined using MTT assay. Briefly, HCT-116、HCT-116 Bax−/− cells were seeded into a 96-well plate at a density of 8,000/well in 96-well plates overnight at a volume of 100 μL medium, the HCT-116、HCT-116 Bax−/− cells treated with the respective agents for the indicated durations for 24h and 48 h. Then exposed to 0.5 mg/ml MTT for 4 h at 37 °C.Then the medium was carefully aspirated and 150 μL of DMSO was added into each well. The cell viability was determined by reduction of MTT after 10-minute incubation at 37°C on a shaking table. Absorbance was measured at 490 nm on a Tecan Sunrise microplate reader (Tecan, Männedorf, Switzerland). The IC50 values were calculated using the relative viability over Bufalin concentration curve.

### Immunoblotting and Immunoprecipitation Analysis

Immunoblotting was performed as described previously [13, 14]. Briefly, 30 μg of protein lysates from blank cells and cells exposed to Bufalin were boiled in SDS-PAGE gel loading buffer, subjected to SDS-PAGE, transferred to PVDF membrane at 100 V for 100 mim at 4°C, and incubated with the respective primary antibody at 4 °C overnight and then incubated with the horseradish peroxidase-conjugated secondary Antibody as described previously [15]to allow detection of the appropriate bands using the chemiluminescence HRP substrate. For immunoprecipitation studies, as we described previously [14]. In brief, Cell lysates were mixed with anti-sirt3, anti-P53 antibody at 4°C overnight. Protein beads A/G were added at a ratio of 1 mg of extract per 30 ml of agarose at 4°C for 3 h. The beads were then pelleted at 3,000 × g for 5 min and washed with lysis buffer five times. The beads boiled in loading buffer and then the lysis were separated on 12% SDS–PAGE pages for immunoblotting.

### Apoptosis assay

The apoptotic rate was determined using Hoechst 33342 staining following fluorescence-microscopic evaluation as well as by annexin V– FITC staining kit (BD Pharmingen, CA, USA) following the manufacturer’s instructions.HCT-116、HCT-116 Bax−/− cells (1×10^5^ cells/well) were cultured in 12‑well flat overnight and subsequently incubated with 1% DMSO (control cells) or 0.05μM,0.1μM,1μM and 2μM Bufalin for 24 h. After fixing in 4% paraformaldehyde (Bio-Rad Laboratories, Inc.) for 15 min at room temperature and staining with 10 ng/ml Hoechst 33342, apoptotic cells were identified by their characteristic nuclear [16]morphology observed under an inverted fluorescence microscope. After treatment with various concentrations (0.05μM,0.1μM,1μM and 2μM) of Bufalin for24 h, HCT-116、HCT-116 Bax−/− cells were harvested, incubated in 500 μL of binding buffer (PH 7.5, 10mM HEPES, 2.5mM CaCl2, and 140mMNaCl) containing Annexin V–FITC and PI for 30min in the dark. Flow cytometry was used to quantify apoptotic of HCT-116 and HCT-116 Bax−/− cells.

### Statistical Analysis

Data are expressed as the mean ± standard deviation (SD) for each experimental group. All the data were analyzed using the SPSS statistical package (SPSS, version 17.0; SPSS, Inc.). Univariate analysis was performed using one-way ANOVA, factorial analysis. The differences between two groups were analyzed, based on the test of least significance difference (LSD). If the variances were not homogenous, mean values were compared using Welch’s test. The differences between two groups were analyzed by Dunnett’sT3. A p value of<0.05 was considered statistically significant.

